# *ahr2*, but not *ahr1a* or *ahr1b*, is required for craniofacial and fin development and TCDD-dependent cardiotoxicity in zebrafish

**DOI:** 10.1101/445213

**Authors:** Jaclyn P Souder, Daniel A Gorelick

## Abstract

The aryl hydrocarbon receptor (AHR) is a ligand-activated transcription factor that binds environmental toxins and regulates gene expression. AHR also regulates developmental processes, like craniofacial development and hematopoiesis, in the absence of environmental exposures. Zebrafish have three paralogues of AHR: *ahr1a*, *ahr1b* and *ahr2*. Adult zebrafish with mutations in *ahr2* exhibited craniofacial and fin defects. However, the degree to which *ahr1a* and *ahr1b* influence *ahr2* signaling and contribute to fin and craniofacial development are not known. We compared morphology of adult *ahr2* mutants and *ahr1a/ahr1b* single and double mutant zebrafish. We found that *ahr1a/ahr1b* single and double mutants were morphologically normal while *ahr2* mutant zebrafish demonstrated fin and craniofacial malformations. At 5 days post fertilization, both *ahr1a/ahr1b* and *ahr2* mutant larvae were normal, suggesting that adult phenotypes are due to defects in maturation or maintenance. AHR was shown to interact with estrogen receptor alpha, yet it is not known whether these interactions are constitutive or dependent on *ahr1* genes. To determine whether estrogen receptors are constitutive cofactors for AHR signaling, we used genetic and pharmacologic techniques to analyze TCDD-dependent toxicity in estrogen receptor and *ahr* mutant embryos. We found that embryos with mutations in *ahr1a/ahr1b* or estrogen receptor genes are susceptible to TCDD toxicity while *ahr2* mutant embryos are TCDD-resistant. Moreover, pharmacologic blockade of nuclear estrogen receptors failed to prevent TCDD toxicity. These findings suggest that *ahr1* genes do not have overlapping functions with *ahr2* in fin and craniofacial development or TCDD-dependent toxicity, and that estrogen receptors are not constitutive partners of *ahr2*.

## INTRODUCTION

The aryl hydrocarbon receptor (AHR) is a ligand-activated transcription factor (Burbach *et al.*, 1992). In the nucleus, AHR forms a heterodimer with aryl hydrocarbon receptor nuclear translocator (ARNT) (Reyes *et al.*, 1992). The AHR-ARNT complex recruits transcription factors and binds xenobiotic response elements DNA sequence (XRE) to regulate gene transcription (Cuthill *et al.*, 1989; Denison *et al.*, 1988a; Denison *et al.*, 1988b; Fujisawa-Sehara *et al.*, 1987; Fujisawa-Sehara *et al.*, 1988; Hapgood *et al.*, 1989). Target genes of AHR include the cytochrome P450 family of genes (e.g. CYP1A), which are involved in the metabolism and clearance of toxic compounds (Burbach, Poland and Bradfield, 1992; Reyes, Reisz-Porszasz and Hankinson, 1992).

The AHR ligand 2,3,7,8-tetrachlorodibenzodioxin (TCDD) is resistant to metabolism and is toxic (Flesch-Janys *et al.*, 1995; Henry *et al.*, 1997). Murine models deficient in AHR (*Ahr-/-*) are resistant to the effects of TCDD, confirming that AHR is required for TCDD-dependent toxicity (Bunger *et al.*, 2008; Bunger *et al.*, 2003; Fernandez-Salguero *et al.*, 1995; Fernandez-Salguero *et al.*, 1996). AHR also binds endogenous ligands to regulate organismal development. Evidence from *Ahr-/-* mice demonstrated that AHR regulates organ development, including hematopoiesis, heart and liver development (Bunger, Glover, Moran, Walisser, Lahvis, Hsu and Bradfield, 2008; Bunger, Moran, Glover, Thomae, Lahvis, Lin and Bradfield, 2003; Fernandez-Salguero, Pineau, Hilbert, McPhail, Susanna, Kimura, Nebert, Rudikoff, Ward and Gonzalez, 1995; Gonzalez and Fernandez-Salguero, 1998; Singh *et al.*, 2011;

Thackaberry *et al.*, 2003). However, the proteins and signaling pathways with which AHR interacts to regulate development are not completely understood. Understanding how AHR functions during development is essential to determine the role of AHR in cases of disrupted development, including congenital heart disease and blood/immune disorders.

To visualize the function of AHR during embryonic development, we turned to the zebrafish model system. Zebrafish produce externally-fertilized embryos that are transparent, allowing rapid analysis of organ formation in live embryos. AHR is conserved throughout the vertebrate lineage, including zebrafish (Hahn *et al.*, 1997). Zebrafish embryos are sensitive to TCDD exposure, exhibiting severe pericardial edema, abnormal heart looping, circulation failure, disrupted erythropoiesis, and reduced jaw length (Belair *et al.*, 2001; Henry, Spitsbergen, Hornung, Abnet and Peterson, 1997; Teraoka *et al.*, 2002). In contrast to mice, zebrafish have three paralogues of *AHR*: *ahr1a*, *ahr1b,* and *ahr2* (Andreasen *et al.*, 2002b; Karchner *et al.*, 2005; Tanguay *et al.*, 1999). In this study, we sought to understand the contribution of each of the zebrafish Ahr receptors to TCDD toxicity, fin development, and craniofacial development.

Zebrafish *ahr2* is thought to be the primary functional homologue of the human AHR based on studies using recombinant Ahr2 protein *in vitro* and *ahr2* mutant zebrafish. Ahr2 binds ligands like TCDD, forms a heterodimer with a zebrafish ARNT paralogue (*arnt2*) on XRE DNA and induces expression of AHR reporter genes like *cyp1a1* (Andreasen *et al.*, 2002a; Andreasen, Spitsbergen, Tanguay, Stegeman, Heideman and Peterson, 2002b; Garcia *et al.*, 2018; Goodale *et al.*, 2012; Prasch *et al.*, 2003; Tanguay, Abnet, Heideman and Peterson, 1999; Tanguay *et al.*, 2000).

Of the zebrafish *ahr* genes, *ahr1a* has the highest sequence similarity to human *AHR.* <u>However</u>, the function of *ahr1a* is not clear. While zebrafish Ahr1a protein was shown to bind Arnt2b at XRE DNA *in vitro*, this interaction was qualitatively weaker than the Ahr2/Arnt2b/XRE interaction (Andreasen, Hahn, Heideman, Peterson and Tanguay, 2002a). Further, Ahr1a was unable to activate XRE reporter activity in cultured cells (Andreasen, Hahn, Heideman, Peterson and Tanguay, 2002a). Ligand binding assays demonstrated that Ahr1a lacks the ability to bind [^3^H]TCDD (Andreasen, Spitsbergen, Tanguay, Stegeman, Heideman and Peterson, 2002b). This suggests that Ahr1a may be a non-functional AHR paralogue. Later studies using morpholino oligonucleotides to knockdown expression of *ahr1a* in zebrafish embryos suggested that *ahr1a* mediates the response to pyrene, polyaromatic hydrocarbon (PAH), and leflunomide (Garner *et al.*, 2013; Goodale, La Du, Bisson, Janszen, Waters and Tanguay, 2012; Incardona *et al.*, 2006). In contrast, a study using TALEN-mediated mutation of *ahr1a* demonstrated that Ahr1a was not required for *cyp1a1* expression (Sugden *et al.*, 2017). Thus, an endogenous role for *ahr1a* has not been elucidated.

In contrast to Ahr1a, zebrafish Ahr1b binds [^3^H]TCDD and Arnt2b and induces XRE-dependent transcription in cultured cells (Karchner, Franks and Hahn, 2005). The EC_50_ of TCDD for Ahr1b-mediated XRE activation is 8-fold higher than for Ahr2 protein (Karchner, Franks and Hahn, 2005). This suggests that Ahr1b may regulate toxic response to dioxins at higher concentrations, while Ahr2 regulates this response at lower concentrations. Morpholino studies suggest that Ahr1b is not required for the response to PAH-mediated toxicity in zebrafish embryos (Garner, Brown and Di Giulio, 2013), but is required in part for leflunomide-induced *cyp1a1* activity (Goodale, La Du, Bisson, Janszen, Waters and Tanguay, 2012). These results, combined with studies investigating *ahr1a* function, suggest that activation of Ahr receptors is ligand-dependent.

We now appreciate that morpholinos are prone to off-target effects (Blum *et al.*, 2015; Gerety and Wilkinson, 2011; Schulte-Merker and Stainier, 2014; Stainier *et al.*, 2015). There is poor correlation between morpholino-induced and mutant phenotypes in zebrafish embryos (Kok *et al.*, 2014). Thus, any observed *ahr1a* and *ahr1b* morpholino phenotypes may not be specific to *ahr1* knockdown. Additionally, genetic compensation could mask a phenotype in *ahr1a* or *ahr1b* single mutants. Here, we generate and analyze single and double *ahr1a* and *ahr1b* mutant embryos.

While ARNT is thought to be an obligate cofactor for AHR (Reyes, Reisz-Porszasz and Hankinson, 1992), additional proteins are also recruited by AHR to mediate changes in gene transcription (Kumar and Perdew, 1999; Nguyen *et al.*, 1999). Several instances of crosstalk between AHR and nuclear estrogen receptors alpha and beta (ERα, ERβ) were described. ERα augmented XRE-dependent transcription in MCF-7 and T47D breast cancer cells and in HuH7 liver cancer cells (Matthews *et al.*, 2005). AHR ligands regulate ER-dependent transcription in cultured cells in an AHR-dependent manner (Kawanishi *et al.*, 2008; Rüegg *et al.*, 2008; Swedenborg *et al.*, 2008). These studies suggest ERs can act as cofactors recruited by AHR. Direct protein-protein interactions between ERs and AHR occured *in vitro* (Beischlag and Perdew, 2005; Matthews, Wihlen, Thomsen and Gustafsson, 2005; Ohtake *et al.*, 2007; Ohtake *et al.*, 2003) and *in vivo* in mouse uterine tissue (Ohtake, Baba, Takada, Okada, Iwasaki, Miki, Takahashi, Kouzmenko, Nohara, Chiba, Fujii-Kuriyama and Kato, 2007; Ohtake, Takeyama, Matsumoto, Kitagawa, Yamamoto, Nohara, Tohyama, Krust, Mimura, Chambon, Yanagisawa, Fujii-Kuriyama and Kato, 2003). These studies demonstrate that nuclear ERs interact with AHR, though it is not well understood whether ERs are constitutive partners of AHR, or if these interactions are ligand- or tissue-dependent.

In the present study, we sought to 1) generate and characterize *ahr1a* and *ahr1b* single and double mutant zebrafish, and 2) determine whether *ahr1* and estrogen receptor genes are required, in addition to *ahr2*, for TCDD-mediated toxicity in zebrafish embryos. Using pharmacologic and genetic approaches, we found that *ahr1* genes are not required for proper fin and craniofacial morphology in adults. Furthermore, neither *ahr1* or estrogen receptor genes are required for TCDD-dependent toxicity in embryos. These results demonstrate that *ahr1* genes are not required for, and do not compensate for, *ahr2* gene function in zebrafish and suggest that ERs are not constitutive partners of AHR.

## MATERIALS AND METHODS

### Zebrafish

Adult zebrafish were raised at 28.5°C on a 14-h light, 10-h dark cycle in the UAB Zebrafish Research Facility and the BCM Zebrafish Research Facility in an Aquaneering recirculating water system (Aquaneering, Inc., San Diego, CA). All zebrafish used for experiments were wild-type AB strain (Westerfield, 2000) and all mutant lines were generated on the AB strain. Zebrafish mutant lines for *esr2a^uab134^*, *esr2b^uab127^* and *gper^uab102^* were previously described (Romano *et al.*, 2017). Zebrafish *esr1^uab119^* mutants were generated in parallel with previously published *esr1^uab118^* mutants using the same guide RNA (Romano, Edwards, Souder, Ryan, Cui and Gorelick, 2017). The *esr1^uab119^* allele harbors an 11-bp deletion resulting in a frameshift mutation at AA165 and premature stop codon at AA168 in the AF-1 domain. All procedures were approved by the UAB and BCM Institutional Animal Care and Use Committees.

### Embryo collection

Adult zebrafish were allowed to spawn naturally in groups. Embryos were collected in intervals of 10 minutes to ensure precise developmental timing and staged following collection, placed in 60 cm^2^ Petri dishes at a density of no more than 100 per dish in E3B media (60X E3B: 17.2g NaCl, 0.76g KCl, 2.9g CaCl_2_-2H_2_O, 2.39g MgSO_4_ dissolved in 1L Milli-Q water; diluted to 1X in 9L Milli-Q water plus 100 μL 0.02% methylene blue), and then stored in an incubator at 28.5°C on a 14-h light, 10-h dark cycle until treatment.

### Embryo treatments

For chemical exposures, embryos were treated in groups of 8-25 in 60 mm^2^ dishes in 7 mL treatment solution beginning at 1 day post fertilization (dpf) until imaging at 3 dpf. Dead embryos were removed daily. Drug stocks were prepared in DMSO and diluted to final concentration in E3B as follows: 2,3,7,8-Tetrachlorodibenzodioxin (TCDD) (10 ng/mL (31 nM), AccuStandard, Inc #D404N); ICI 182,780 (Fulvestrant) (10 μM, Sigma #I4409). DMSO was used as vehicle control (0.1%). The concentration of TCDD was chosen as 100% of wild-type embryos demonstrated cardiotoxicity at 3 dpf.

### CRISPR-Cas9 Mutant Generation

Cas9 mRNA and gRNAs for *ahr1a, ahr1b,* and *ahr2* mutants were generated as previously described for *esr1, esr2a, esr2b,* and *gper* mutants (Romano, Edwards, Souder, Ryan, Cui and Gorelick, 2017). Oligonucleotides containing target site sequences are shown in Table 1. Briefly, oligos were hybridized to generate double-stranded target DNA and annealed into digested pT7-gRNA, then gRNAs were synthesized and purified. Cas9 mRNA was generated using a linearized pT3TS-nCas9n plasmid, transcribed, and purified.

**Table 1.**
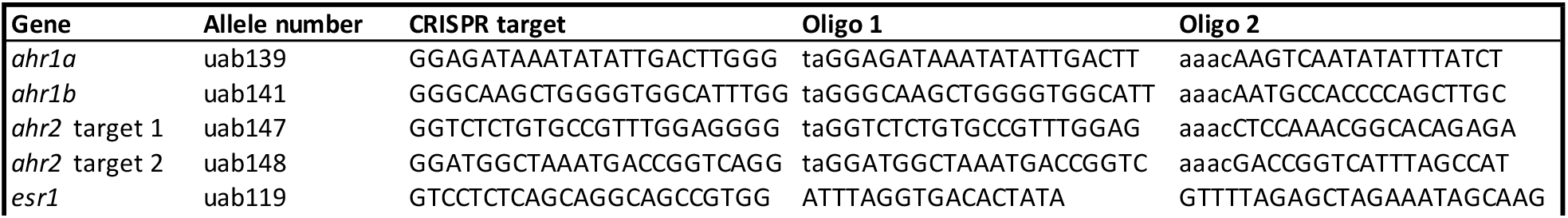
gRNA targets and primers for mutant generation.

For mutant generation, one-cell-stage embryos were injected using glass needles pulled on a Sutter Instruments Fleming/Brown Micropipette Puller, model P-97 and a regulated air-pressure micro-injector (Harvard Apparatus, NY, PL1–90). Each embryo was injected with a 1 nl solution of 150 ng/μl of Cas9 mRNA, 30 ng/μl of gRNA per target (for a maximum of two targets per injection) and 0.1% phenol red. Mixtures were injected into the yolk of each embryo. At least 100 injected embryos per gRNA pair were raised to adulthood and crossed to wild-type fish (AB strain) to generate F1 embryos. F1 offspring with heritable mutations were sequenced to identify loss of function mutations.

### Genotyping

Genomic DNA was isolated from individual embryos or tail biopsies from individual adult fish by incubation in 50 μL ELB (10 mM Tris pH 8.3, 50 mM KCl, 0.3% Tween 20) with 0.5 μL proteinase K (800 U/ml, NEB) in 96 well plates, one sample per well, at 55°C for 8 hours. Proteinase K was inactivated by incubation at 98°C for 10 minutes and DNA was stored at −20°C. Genotyping was performed by PCR and high-resolution melting curve analysis as described (Parant *et al.*, 2009; Romano, Edwards, Souder, Ryan, Cui and Gorelick, 2017). All melting curves were generated with a Bio-Rad CFX96 Real-Time System over a 70-95°C range and analyzed with Bio-Rad CFX Manager 3.1 software. All mutations were confirmed by TA cloning and sequencing.

### Embryo and Larval Zebrafish Imaging

All embryos were imaged with a Nikon SMZ25 microscope equipped with a Hamamatsu ORCA-Flash4.0 digital CMOS camera. Images were equally adjusted for brightness and contrast in Adobe Photoshop CC 2018. For imaging of embryo fin and jaw, embryos were fixed at 5 dpf overnight in 4% formaldehyde in phosphate-buffered saline (PBS) and stored in 70% ethanol at −20°C until imaging. Embryos were embedded in 3% methyl cellulose for imaging. Fin length was measured using ImageJ (Schneider *et al.*, 2012) by free-tracing along the edge of each fin and taking the average of left and right fin length for a minimum of n=8 fish per genotype. Jaw length was measured in Nikon AR elements software by measuring from the anterior edge of the eyes to the tip of the jaw line as previously described (Teraoka, Dong, Ogawa, Tsukiyama, Okuhara, Niiyama, Ueno, Peterson and Hiraga, 2002) for a minimum of n=7 fish per genotype. Embryos from heterozygous parents were imaged as a mixed clutch and genotyped following imaging. Eye diameter was measured to control for differences in larval size at time of fixation (Parichy *et al.*, 2009). Fin and jaw length measurements from each larva were adjusted for size with a normalization factor created by dividing the mean eye diameter of all embryos by the eye diameter of each individual embryo. Measurements for each embryo were multiplied by this normalization factor for statistical analysis. For imaging of embryos following TCDD treatment, embryos were imaged in E3B in 60 mm^2^ dishes in 0.02% tricaine anesthetic. For embryos treated as mixed clutch, embryos were genotyped following imaging.

### Adult zebrafish imaging

Anal and caudal fins from adult zebrafish (8-12 months old) were imaged via brightfield microscopy with a Leica M205 FA microscope equipped with a Leica DFC3000 G camera under 0.01% tricaine anesthesia. Pectoral fins from adults (6-7 months old) were captured from freely swimming, unanesthetized adult zebrafish with a Canon EOS Rebel T5 digital camera and adjusted for brightness and contrast in Adobe Lightroom CC.

### Micro-CT Analysis

Adult zebrafish heads (4 per sex per genotype) were collected at 3-6 months post-fertilization and fixed overnight (18-24 hours) in 4% formaldehyde in PBS and stored in 70% ethanol at −20°C until further analysis. For imaging, heads were embedded in 1.5-2 mL microcentrifuge tubes (Fisherbrand, #05-408-129 and Genesee, #24-283) in 1% agarose. Scanning was performed on a Bruker SkyScan 1272 Scanner at 50.0 kV/200 μA with 9 μm voxel resolution and 280 ms exposure time. 3D reconstructions were made with CTVox software and relative skull and jaw lengths were measured in ImageJ.

### Experimental Design and Data Analysis

TCDD exposure experiments were performed on at least 8 embryos from a single clutch per treatment group or vehicle control group. Experiments were performed on 2-5 clutches for each genotype except for *ahr1a^uab139/-^*, which was performed on one clutch of 24 embryos per treatment group. For *esr2b^uab127^*, embryos were derived from breeding heterozygous parents and analyzed as a mixed clutch of embryos (*esr2b^uab127-/-,-/+,+/+^*) where experimenter was blind to genotype. One *esr2b* mutant clutch was genotyped to confirm presence of homozygous and heterozygous embryos in Mendelian ratios. Other clutches are presumed to contain 25% homozygous mutants. Mean larval fin length and jaw length were measured from at least 7 larvae per genotype. Values were adjusted for eye diameter to control for differences in growth between larvae. Genotypes were compared using one-way ANOVA with Tukey’s multiple comparisons test. Mean ventral skull width and standard length of adult zebrafish was used for comparing genotypes by sex using a two-way ANOVA with Tukey’s multiple comparisons test where factor 1 is genotype and factor 2 is sex. Statistical significance was set at p ≤ 0.05. GraphPad Prism 7.0d software was used for all statistical analyses and for producing graphs.

## RESULTS

### Generation of aryl hydrocarbon receptor mutant zebrafish

To compare the function of the three zebrafish aryl hydrocarbon receptors (*ahr1a, ahr1b* and *ahr2*), we utilized CRISPR-Cas9 technology to generate single and double *ahr1* mutants (Figure 1A-B, Table 1: *ahr1a^uab139^*, *ahr1b^uab141^*) and an *ahr2* mutant (Figure 1C, Table 1: *ahr2^uab147;uab148^*). For *ahr1a*, we generated embryos with a 12-basepair (bp) deletion and 7-bp insertion within the transactivation domain in exon 10, which results in a frameshift mutation at amino acid (AA) 440 and a premature stop codon at AA446.

**Figure 1.**
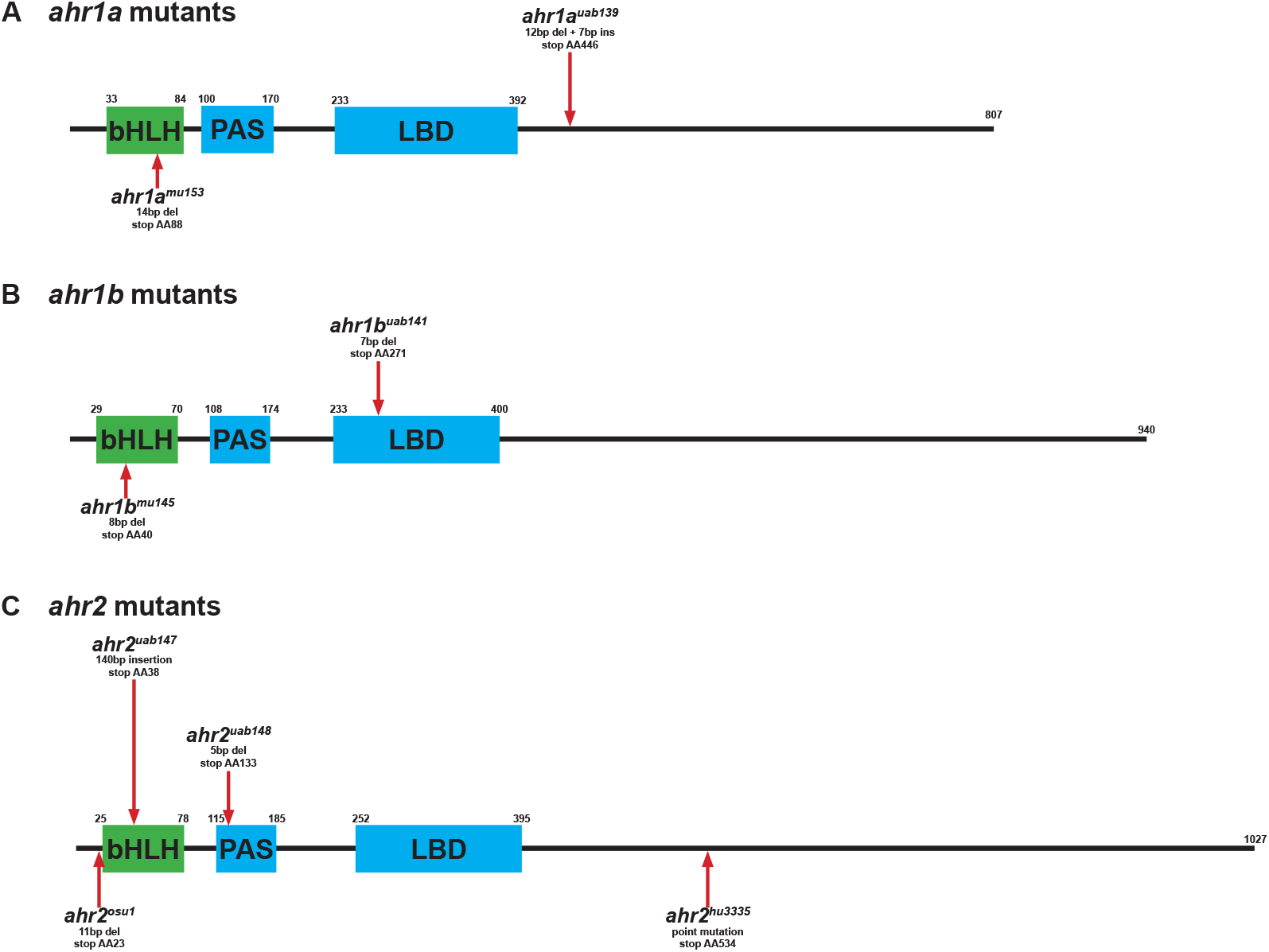
Zebrafish aryl hydrocarbon receptor mutants. The amino acid sequence for the zebrafish aryl hydrocarbon receptors 1a **(A)**, 1b **(B)**, and 2 **(C)** are represented by the black line. Protein domains basic helix-loop-helix (bHLH), per-arnt-sim (PAS) and ligand binding domain (LBD) are represented by green and blue boxes. Numbers indiciate the starting and ending point of these domains in the amino acid sequence. Mutations created in each of these receptors are represented by red arrows pointing to location of the premature stop codon created by the mutation. Mutants generated for the present study are located above each gene, while previously published mutants are located below. bp, DNA basepairs; del, deletion; ins, insertion; AA, amino acid. *mu145*, *mu153* from Sugden et al., 2017; *osu1* fromGarcia et al., 2018; *hu3335* from Goodale et al., 2012.

For *ahr1b*, we generated embryos with a 7-bp deletion in exon 7, resulting in a frameshift mutation at AA255 and premature stop codon within the ligand binding domain at AA271. These guide RNAs were injected simultaneously to generate both single *ahr1a^uab139^* and *ahr1b^uab141^* mutants as well as a double *ahr1a^uab139^/ahr1b^uab141^* mutant. Finally, we generated embryos with two different mutations in *ahr2*: uab147 and uab148. Guide RNAs to two targets were created: target 1 in exon 2 of the basic helix-loop-helix (bHLH) domain and target 2 in exon 3 upstream of the PAS-A domain. Guide RNAs for each of these targets were injected simultaneously for uab147 (target 1) and uab148 (target 2). The *ahr2^uab147^* allele contains a 140-bp insertion in exon 2, resulting in a frameshift mutation at AA33 and premature stop codon at AA38 in the bHLH domain. Embryos with this mutation also inherited the mutation for uab148 (5-bp deletion in exon 3 resulting in a frameshift mutation at AA98 and premature stop codon at AA133), although this mutation is downstream of uab147 and therefore does not influence the mutant transcript. The predicted effect of each of these mutations for *ahr1a*, *ahr1b*, and *ahr2* is loss of functional protein. Embryos from all mutants were grossly morphologically normal and demonstrated normal mendelian ratios from heterozygous crosses. Previously published zebrafish *ahr* mutants are highlighted in Figure 1 for comparison (Garcia, Bugel, Truong, Spagnoli and Tanguay, 2018; Goodale, La Du, Bisson, Janszen, Waters and Tanguay, 2012; Sugden, Leonardo-Mendonça, Acuña-Castroviejo and Siekmann, 2017).

### Ahr2, but not Ahr1a or Ahr1b, is required for proper adult fin morphology

Previous studies have determined that *ahr2* is important for proper fin morphology in adult zebrafish (Garcia, Bugel, Truong, Spagnoli and Tanguay, 2018; Goodale, La Du, Bisson, Janszen, Waters and Tanguay, 2012). We therefore first validated our *ahr2* mutant line by analyzing fin morphology of adult zebrafish mutants. We found that *ahr2^uab147/-^* mutant adults of both sexes (n=8 per sex) had severely deformed or absent anal, caudal, and pectoral fins (Figure 2E-E’). To test whether this phenotype was specifically due to loss of *ahr2* function, we analyzed fin morphology in *ahr1a^uab139/-^*, *ahr1b^uab141/-^*, and *ahr1a^uab139/-^*;*ahr1b^uab141/-^* mutant adult zebrafish (n=8 per sex per genotype). We found that fins developed normally in these mutants in males and females (Figure 2A-D), suggesting that Ahr2 is responsible for proper fin morphology independently of Ahr1a and Ahr1b.

**Figure 2.**
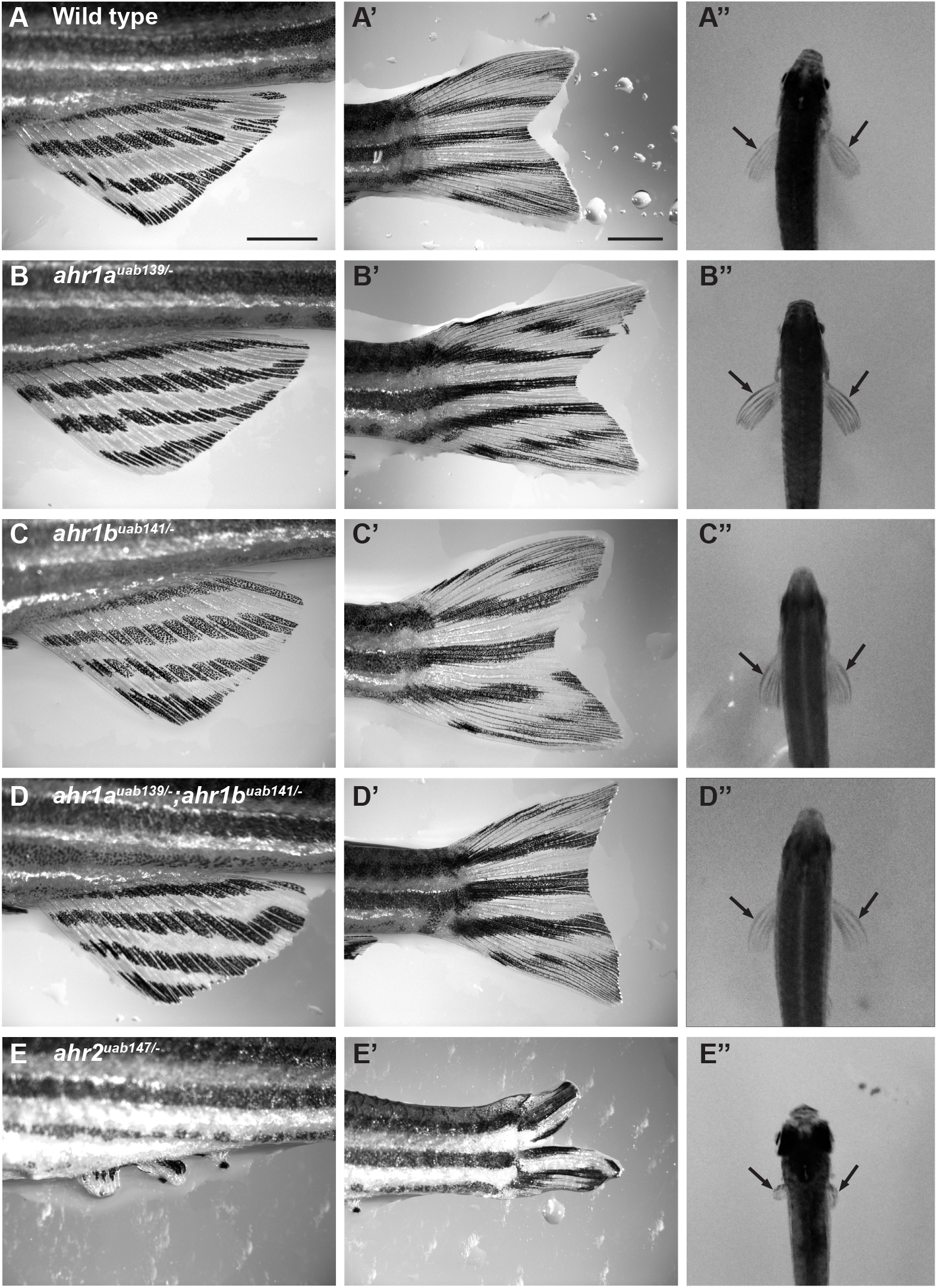
*ahr2*, but not *ahr1* mutants, have abnormal fin morphology. **(A-E, A’-E’)**Images of adult male zebrafish with genotype indicated. (A-E), images of the anal fin, (A’-E’), caudal fin. **(A”-E”),** pectoral fins (arrows). Absence of fins and fin deformities are present in anal, caudal, and pectoral fins in *ahr2* -/-mutant fish (E, E’), but not in *ahr1* mutants or wild-type fish (A-D, A’-D’). Scale bars in panels A and A’ = 2mm and apply to B-E and B’-E’, respectively.

### Ahr2, but not Ahr1a or Ahr1b, is required for proper adult craniofacial morphology

Previous studies suggested that *ahr2* is important for proper craniofacial morphology (Garcia, Bugel, Truong, Spagnoli and Tanguay, 2018). To explore gross changes in craniofacial morphology in our *ahr2* mutants, we performed μCT analysis of adult zebrafish heads at 3-6 months of age. We found that *ahr2* mutant adults (n=4 per sex) have a 29% or 38% decreased ventral skull width (in males and females, respectively) as measured by the ratio of the width between the paired opercular bones and paired dentate bones (Figure 3A-D; male: wildtype=2.67±0.11 (mean±SD), *ahr2^147/-^* =1.89±0.09, mean difference=0.78, p<0.0001; female: wildtype=2.85±0.23, *ahr2^147/-^* =1.90±0.24, mean difference=0.95, p<0.0001; two-way ANOVA). This ratio measures the width of the broadest part of the caudal skull to the narrowest part of the cranial skull with easily identifiable landmarks on 3D reconstructions from μCT data (Figure 3G). To determine if *ahr1* genes are also important for proper craniofacial morphology, we performed μCT analysis of *ahr1a/ahr1b* double mutant fish. We found no significant difference between *ahr1a/ahr1b* double mutant zebrafish and wildtype (Figure 3E-F; *ahr1a^139/-^;ahr1b^141/-^* male=2.47±0.16 (mean±SD), mean difference=0.20, p=0.21; *ahr1a^139/-^;ahr1b^141/-^* female=2.73±0.04, mean difference=0.12, p=0.56; two-way ANOVA), while there was a significant difference between *ahr1a/ahr1b* double mutants and *ahr2* mutants (male: p=0.0003, female: p<0.0001).

**Figure 3.**
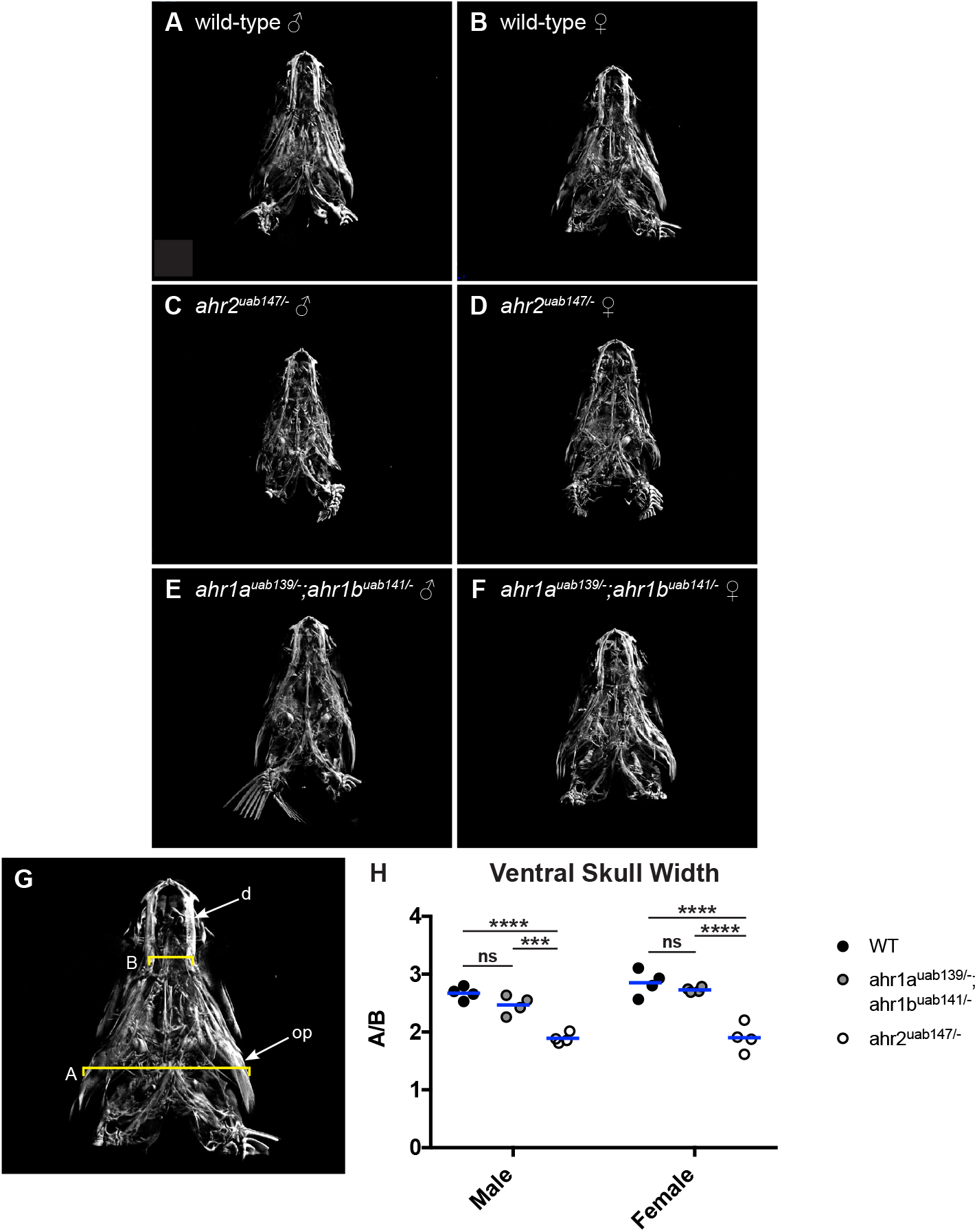
*ahr2* but not *ahr1* mutant adults have reduced ventral skull width. **(A-F)** Micro-CT images of male (A,C,E) and female (B,D,F) adult zebrafish heads of indicated genotype **(G)** Measurements are indicated on the wild-type female from (B). d=dentate bone, op=opercle, A=width between paired opercle bones, B=width between paired dentate bones. **(H)** Male and female *ahr1* mutants do not have significantly different skull width compared to wild-type by A/B ratio, while *ahr2* male and female mutants have a significantly decreased A/B ratio compared to wild-type and *ahr1a/ ahr1b* double mutants. ^***^p<0.001,^****^p<0.0001, ns not significant (p>0.05), two-way ANOVA with Tukey’s multiple comparisons test.

It is possible that the more narrow skulls observed in *ahr2* mutants is due to a general growth defect. To test for a growth defect, we analyzed standard length (Parichy, Elizondo, Mills, Gordon and Engeszer, 2009), an established parameter of growth and development. We found no significant difference in standard length between wild type and mutants of the same sex (Table 2). These results suggest that Ahr2 regulates craniofacial morphology independently of Ahr1a and Ahr1b.

**Table 2.** Standard length (SL) of adult zebrafish used for μCT analysis. Fish are grouped by sex and genotype (n=4 individuals per group). Mean and standard deviation (SD) are reported in mm. *p* value describes comparison between mutant and wild-type fish of the same sex. Two-way ANOVA with Tukey’s multiple comparisons.

| Genotype | Sex | SL (mm) |  |  | *p* |
| --- | --- | --- | --- | --- | --- |
|  |  | mean | SD | n |  |
| Wild-type | Male | 25.87 | 1.23 | 4 | - |
|  | Female | 27.2 | 2.79 | 4 | - |
| *ahr1a*<sup>*uab139/-*</sup>;*ahr1b*<sup>*uab141/-*</sup> | Male | 26.64 | 1.25 | 4 | 0.84 |
|  | Female | 29.56 | 1.93 | 4 | 0.23 |
| *ahr2*<sup>*uab147/-*</sup> | Male | 22.47 | 1.8 | 4 | 0.06 |
|  | Female | 25.42 | 2.29 | 4 | 0.42 |
| **Table 2. Standard length (SL) of adult zebrafish used for μCT analysis.** Fish are grouped by sex and genotype (n=4 individuals per group). Mean and standard deviation (SD) are reported in mm. *p* value describes comparison between mutant and wild-type fish of the same sex. Two-way ANOVA with Tukey's multiple comparisons. |  |  |  |  |  |

### Aryl hydrocarbon receptors are not required for larval fin growth

Since we observed fin defects in *ahr2* mutant adult zebrafish, we tested whether fin defects were present in larval development. We compared the length of the pectoral fins at 5 dpf in *ahr2*, *ahr1a*, *ahr1b*, and *ahr1a/ahr1b* mutant and wild-type larvae. All measurements were corrected for embryo size using eye diameter (Table 3). We found no significant difference in pectoral fin length between *ahr2* mutant and wildtype larvae (Table 3, Figure 4 A,E,G; wildtype=472.9±9.0 μm (mean±SD), *ahr2^uab147/-^*=464.0±29.49 μm, p=0.85, one-way ANOVA). We next compared pectoral fin length between *ahr1* mutants and wild-type larvae, and saw no significant difference (Table 3, Figure 4A-D,G; *ahr1a^uab139/-^*=461.4±22.2 μm, p=0.68; *ahr1b^uab141/-^*=497.4±18.29 μm, p=0.06; *ahr1a^uab139/-^;ahr1b^uab141/-^*=471.1±23.21 μm, p=0.9997). These results suggest that aryl hydrocarbon receptors are not required for embryonic fin development.

**Figure 4.**
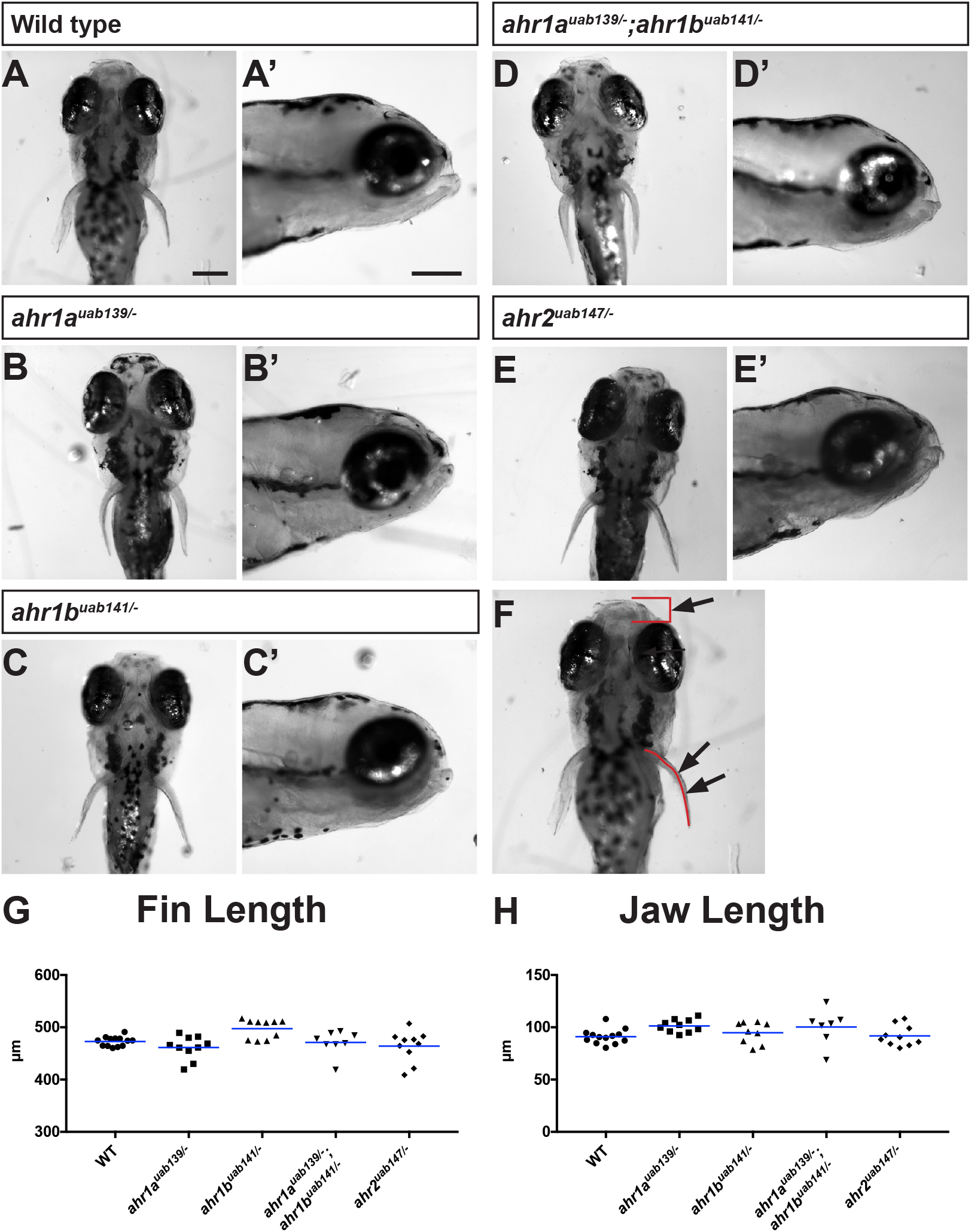
*ahr1* and *ahr2* mutant larvae have normal fin and jaw morphology. **(A-E)**Represenative images of jaw and fin morphology for wild-type, *ahr1*, and *ahr2* mutant larvae at 5 days post fertilization. (A-E) Embryo head and pectoral fins, ventral view. (A’-E’) Jaw profile, sagittal view. **(F)** Example measurements used for data in graphs quantifying fin **(G)** and jaw **(H)** length. Arrows point to regions measured for fin and jaw length quantification outlined in red. There is no significant difference in fin length or jaw length in *ahr1* or *ahr2* mutant larvae compared to wild-type. p>0.05, one-way ANOVA with Tukey’s multiple comparisons test. Scale bars in A and A’ = 200 μm and apply to panels B-E and B’-E’, respectively. Fin and jaw lengths were corrected for standard length of each fish using eye diameter.

**Table 3.**
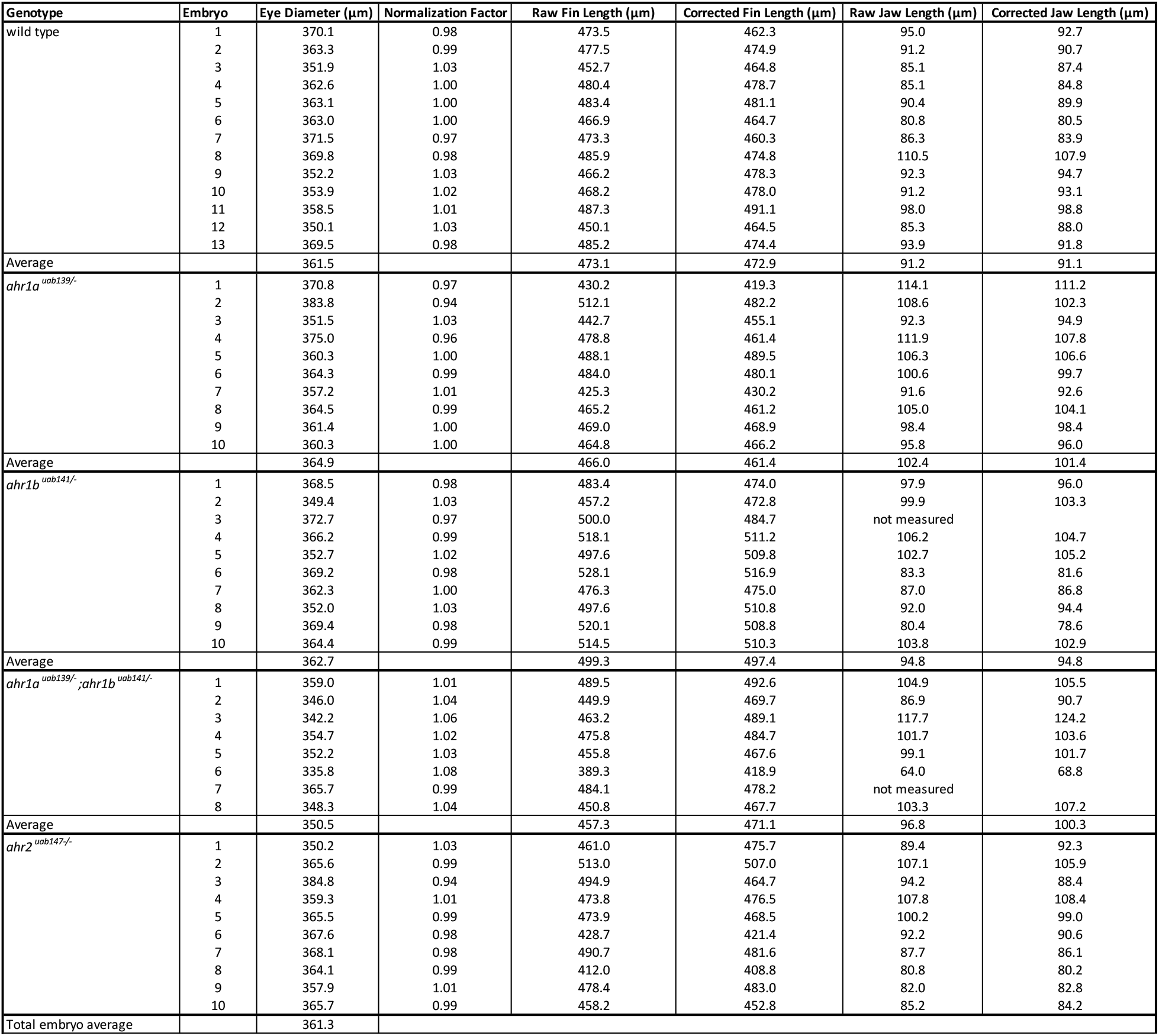
Embryo fin and jaw length normalized to eye diameter. Embryos are grouped by genotype with 7-13 embryos per group. All embryos were fixed at 5 days post fertilization prior to imaging. The eye diameter for each embryo was measured, then the average eye diameter of all embryos was calculated ("Total embryo average"). To create a normalization factor for each embryo, the total embryo average eye diameter was divided by the individual embryo eye diameter. The normalization factor was then multiplied by the raw fin length or raw jaw length values to create corrected values for each measurement. Corrected values were used in all statistical analyses.

### Aryl hydrocarbon receptors are not required for larval jaw growth

Since we observed changes in craniofacial morphology in *ahr2* mutants, we also compared the craniofacial morphology of *ahr1a/ahr1b and ahr2* mutant and wild-type larvae at 5 dpf. Since the skull is mostly cartilaginous at this developmental stage and opercular bones are not fully formed, we instead compared morphology by measuring the length of the jaw from the anterior tip of the eye to the end of the jaw, as has been previously used to compare jaw growth in embryos (Teraoka, Dong, Ogawa, Tsukiyama, Okuhara, Niiyama, Ueno, Peterson and Hiraga, 2002). All values were corrected for larval size using eye diameter (Table 3). When comparing jaw length between *ahr2^uab147/-^* mutant and wildtype larvae, we found no significant difference (Table 3, Figure 4A,E,H; wildtype=91.09±7.02 μm (mean±SD), *ahr2^uab147/-^*=91.78±9.71 μm, p=0.9998, one-way ANOVA). We next compared jaw length between *ahr1* mutant and wildtype larvae, and did not observe a significant difference compared to wildtype in single or double *ahr1* mutants (Table 3, Figure 4A-D,H; *ahr1a^uab139/-^*=101.4±6.08 μm, p=0.12; *ahr1b^uab141/-^*=94.83±10.29 μm, p=0.91; *ahr1a^uab139/-^;ahr1b^uab141/-^*=100.3±17.04 μm, p=0.30). These results suggest that aryl hydrocarbon receptors are not required for embryonic jaw growth.

### Ahr2, but not Ahr1a or Ahr1b, is required for TCDD-induced cardiotoxicity in embryos

2,3,7,8-tetrachlorodibenzodioxin (TCDD) is a persistent environmental toxin that acts as a carcinogen in humans and causes cardiotoxicity and mortality at nanomolar concentrations in zebrafish embryos (Belair, Peterson and Heideman, 2001; Flesch-Janys, Berger, Gurn, Manz, Nagel, Waltsgott and Dwyer, 1995; Henry, Spitsbergen, Hornung, Abnet and Peterson, 1997). Ahr2 is known to mediate the toxic effects of TCDD (Garcia, Bugel, Truong, Spagnoli and Tanguay, 2018; Goodale, La Du, Bisson, Janszen, Waters and Tanguay, 2012; Tanguay, Abnet, Heideman and Peterson, 1999), yet it is not clear whether zebrafish *ahr1* genes are also required for TCDD toxicity. To test this, we exposed zebrafish with mutations in each of the aryl hydrocarbon receptors (*ahr2, ahr1a, ahr1b*) and *ahr1a/ahr1b* double mutants to TCDD. 100% of *ahr1a* and *ahr1b* single and double mutants (1-3 clutches per genotype, 3-25 embryos per clutch) demonstrated severe pericardial edema and abnormal heart looping, which are characteristic of TCDD toxicity in wildtype embryos at 3 dpf (Table 4, Figure 5A-H). In contrast, 100% of *ahr2* homozygous mutants (2 clutches, 3-4 embryos per clutch) were resistant to TCDD toxicity (Table 4, Figure 5I-J). 100% of *ahr2* -/+ and +/+ embryos from the same clutches (2 clutches, 3-12 embryos per clutch) also demonstrated TCDD toxicity or mortality at 3 dpf, suggesting a single copy of *ahr2* is sufficient to induce TCDD toxicity (Table 4, Figure 5K-L). Less than 18% mortality was observed in all DMSO-treated embryos from the same clutches and ≤17% of DMSO-treated embryos exhibited pericardial edema or abnormal heart looping at 3 dpf (Table 4), providing evidence that observed cardiotoxicity is specific to TCDD treatment and not a side effect of mutation. Overall, these results suggest that Ahr1a and Ahr1b are not required for TCDD toxicity.

**Figure 5.**
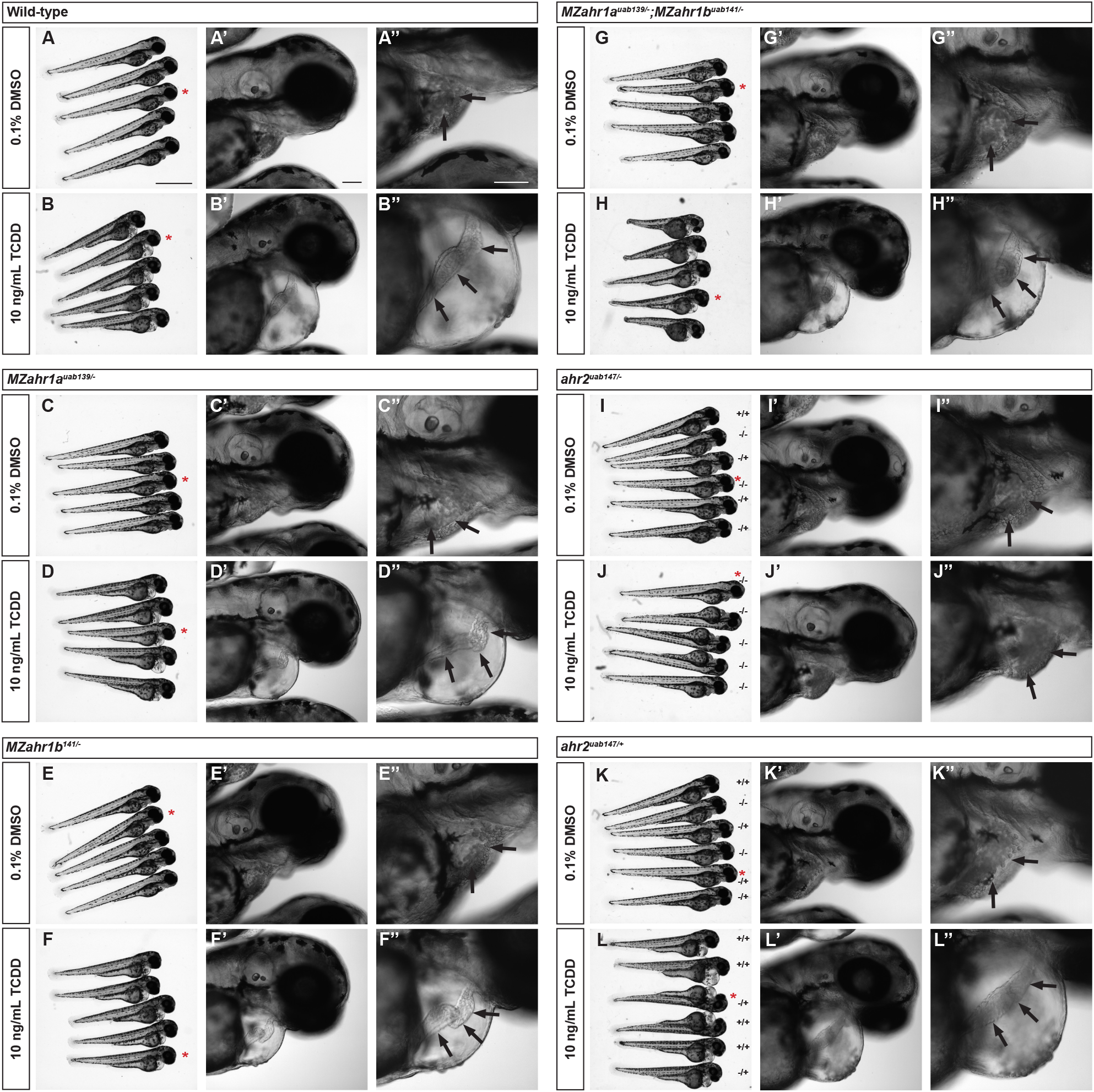
*ahr2*, but not *ahr1* mutants, are responsive to TCDD. **(A-L)**Representative images of embryos at 3 days post fertilization exposed to DMSO or TCDD. Embryos derived from heterozygous parents were genotyped following imaging (denoted as wild-type +/+, heterozygous -/+, or homozygous -/-, panels I-L). Embryos derived from homozygous parents are denoted as maternal zygotic (MZ). Images in panels A-L display overview images of treated embryos and red asterisks denote embryos detailed in panels A’-L’ and A’’-L’’. Black arrows in panels A’’-L’’ point to the heart within the pericardial cavity. Obvious pericardial edema and failure of heart looping is present in 100% of embryos from all TCDD-treated groups with the exception of *ahr2* -/- mutants (J, J’, J’’), in which 0% of embryos displayed cardiotoxicity. Scale bar in panel A = 1000 μm and applies to panels B-L. Scale bars in panels A’ and A’’ = 100 μm and apply to B’-L’ and B’’-L’’, respectively.

**Table 4.**
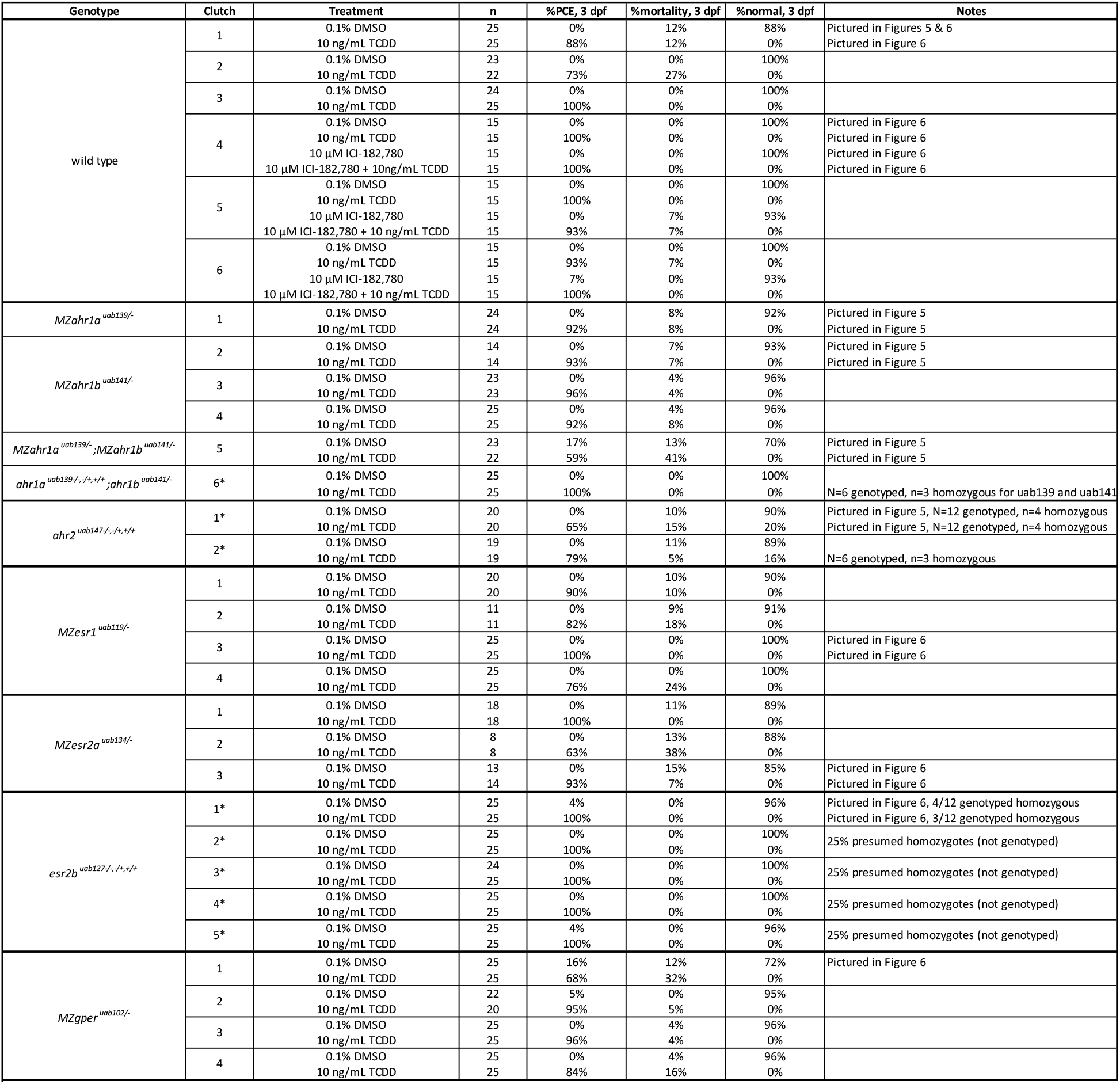
TCDD toxicity in *ahr1, ahr2*, and estrogen receptor mutant embryos. Embryos are grouped by genotype and clutch, with each clutch of embryos split between two treatment groups: 0.1% DMSO (vehicle) or 10 ng/mL TCDD. An additional two treatment groups are present for pharmacologic inhibition of estrogen receptors in wild-type embryos (10 μM ICI182,780 and 10 μM ICI182,780 + 10 ng/mL TCDD). Asterisk (*) denotes clutches derived from heterozygous parents and thus containing homozygous, heterozygous and wild type siblings. All genotypes included in the clutch are listed in the "Genotype" column. The number of embryos genotyped and number of homozygous embryos are listed in the "Notes" column. Mutant embryos derived from homozygous parents are denoted maternal zygotic (MZ). The percent of embryos with pericardial edema (PCE), percent mortality, and percent normal embryos at 3 days post fertilization (dpf) are listed in right three columns. All TCDD-treated groups have 0% normal embryos at 3 dpf, except for mixed-genotype *ahr2* mutant clutches. All normal TCDD-exposed embryos are ahr2 ^*uab147/-*^. The "Notes" column denotes embryos chosen as representative images in Figures 5 & 6, with number of homozygous embryos listed for mixed genotype clutches.

### Estrogen receptors are not required for TCDD-induced cardiotoxicity in embryos

AHR is a ligand-dependent transcription factor that partners with the aryl hydrocarbon receptor nuclear translocator (ARNT) along with other cofactors at dioxin response elements in the DNA to mediate changes in gene transcription (Burbach, Poland and Bradfield, 1992; Reyes, Reisz-Porszasz and Hankinson, 1992). Estrogen receptors alpha and beta (ERα and ERβ) have been shown to directly interact with AHR *in vitro* and *in vivo* to regulate gene expression (Beischlag and Perdew, 2005; Ohtake, Baba, Takada, Okada, Iwasaki, Miki, Takahashi, Kouzmenko, Nohara, Chiba, Fujii-Kuriyama and Kato, 2007; Ohtake, Takeyama, Matsumoto, Kitagawa, Yamamoto, Nohara, Tohyama, Krust, Mimura, Chambon, Yanagisawa, Fujii-Kuriyama and Kato, 2003), yet it is unclear whether this interaction is constitutive or conditional. To test the hypothesis that ERs are recruited by AHR2 following TCDD treatment, we exposed zebrafish with mutations in each of the nuclear estrogen receptors *esr1* (ERα)*, esr2a* (ERβ1) and *esr2b* (ERβ2) to TCDD. If ERs are required for TCDD toxicity, then ER mutant embryos should be resistant to TCDD toxicity, similar to *ahr2* mutant embryos. We found, however, that 100% of ER mutant embryos (3-5 clutches per genotype, 3-25 embryos per clutch) displayed abnormal pericardial edema and heart looping at 3 dpf following TCDD treatment (Table 4, Figure 6A-H).

**Figure 6.**
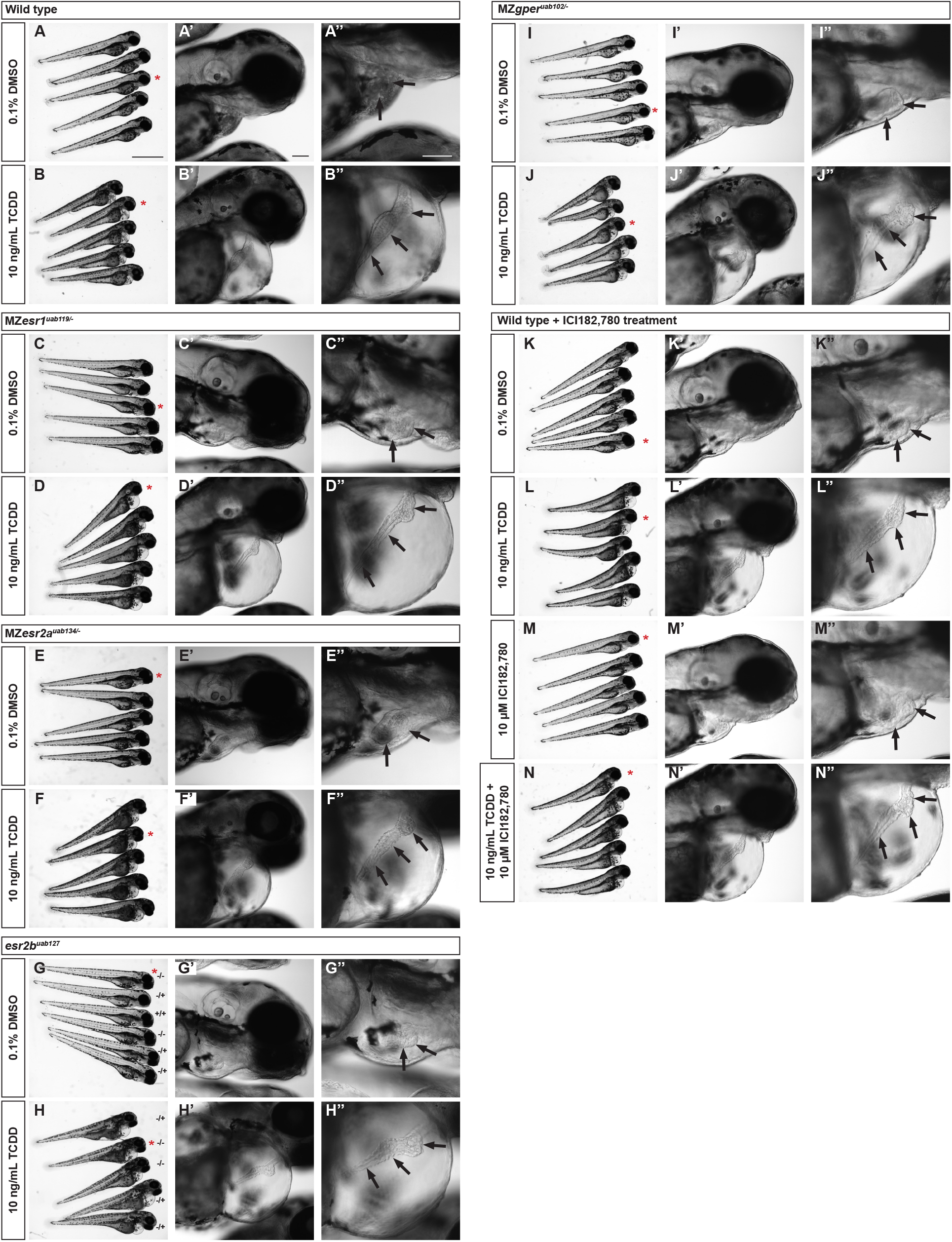
Estrogen receptors are not required for TCDD toxicity. **(A-N)** Embryos of the indicated genotype were exposed to DMSO, TCDD, or the estrogen receptor antagonist ICI182,780 at 3 dpf. *esr2b* mutant embryos were derived from heterozygous parents, thus the clutch contained a mix of wild type and mutant embryos (denoted as wild-type +/+, heterozygous -/+, or homozygous -/-, panels G-H). Other mutant embryos were derived from homozygous parents and are denoted as maternal zygotic (MZ). Images in panels A-N display overview images of treated embryos and red asterisks denote embryos detailed in panels A’-N’ and A’’-N’’. Black arrows in panels A’’-N’’ point to the heart within the pericardial cavity. Pericardial edema and failure of heart looping is present in 100% of embryos from all TCDD-treated groups. Scale bar in panel A = 1 mm and applies to panels B-N. Scale bars in panels A’ and A’’ = 100 μm and apply to B’-N’ and B’’-N’’, respectively. Wild-type embryos in panels A-B are the same embryos shown in Figure 5A-B.

It is possible that there is a compensatory relationship between Esr1, Esr2a, and Esr2b as cofactors for Ahr2 (e.g. *esr2b* is upregulated in *esr1* mutants to mediate the effects of TCDD). To test this hypothesis, we treated embryos with TCDD together with the pan-nuclear estrogen receptor antagonist, ICI-182,780 (Fulvestrant). We saw that pharmacological inhibition of ERs did not protect embryos from TCDD toxicity, as 100% of co-treated embryos (3 clutches, 15 embryos per clutch) demonstrated pericardial edema and abnormal heart looping or mortality at 3 dpf (Table 4, Figure 5I-N). To further investigate the hypothesis that estrogen receptors are required for *ahr2* activity, we tested whether the G protein-coupled estrogen receptor (*gper*) is required for TCDD toxicity. In agreement with results from nuclear ERs, 100% of *gper* mutants (4 clutches, 20-25 embryos per clutch) demonstrated characteristic TCDD toxicity or mortality at 3 dpf (Table 4, Figure 5I-J). Less than ≤13% mortality is observed in all DMSO- or ICI-182,780-treated embryos from the same clutches and ≤16% of embryos demonstrate PCE or abnormal heart looping at 3 dpf (Table 4), providing evidence that observed cardiotoxicity is specific to TCDD treatment and not a side effect of mutation or ICI182,780 treatment. These results suggest that ERs are not required for TCDD toxicity, and that estrogen receptors are not required cofactors for Ahr2.

## DISCUSSION

We find that *ahr2* regulates TCDD-toxicity and fin and craniofacial morphology in zebrafish, consistent with previous results (Garcia, Bugel, Truong, Spagnoli and Tanguay, 2018; Goodale, La Du, Bisson, Janszen, Waters and Tanguay, 2012; Prasch, Teraoka, Carney, Dong, Hiraga, Stegeman, Heideman and Peterson, 2003). Using new *ahr1* mutants, we demonstrate that these functions of *ahr2* are independent of *ahr1* genes. Further, we provide evidence that Ahr2 recruits cofactors in a context-dependent manner.

The fin deformities observed in adult *ahr2^uab147/-^* mutants were severe, resulting in dysmorphic caudal and pectoral fins and absent anal fins in adult zebrafish of both sexes (Figure 2). This defect first presents in juvenile stages of growth, as mutant larvae have normal fins at 5 dpf (Figure 4). Previous studies reported abnormal morphology in adult fins of *ahr2* mutant fish (Garcia, Bugel, Truong, Spagnoli and Tanguay, 2018; Goodale, La Du, Bisson, Janszen, Waters and Tanguay, 2012), resulting in a “damaged fin” phenotype. However, in our mutant we observe almost complete absence of anal fins, more akin to the zebrafish mutant *finless* (Haffter *et al.*, 1996; van Eeden *et al.*, 1996). This mutant was later discovered to harbor a mutation in the *edar* gene encoding the ectodysplasin A receptor (Harris *et al.*, 2008). Similar to *ahr2^uab147/-^* mutants, *edar* homozygous mutants demonstrate normal larval fin development, but have an absence of all fins (pectoral, anal, caudal, pelvic, and dorsal) in adulthood. *edar* mutants first demonstrate fin defects in juvenile metamorphosis and adulthood. One hypothesis is that Ahr2 is involved in Edar signaling to regulate fin development. This is supported by the fact that the ectodysplasin A receptor and its ligand ectodysplasin are structurally analogous to tumor necrosis factor receptor (TNFR), and its ligand, tumor necrosis factor alpha (TNFα), a regulator of inflammation. AHR has been shown to regulate inflammation in a ligand-dependent manner (Quintana *et al.*, 2008). Further, *Ahr* null mice have altered populations of immune cells (Fernandez-Salguero, Pineau, Hilbert, McPhail, Susanna, Kimura, Nebert, Rudikoff, Ward and Gonzalez, 1995; Singh, Garrett, Casado and Gasiewicz, 2011). It is possible that an endogenous ligand for Ahr2 activates the Edar pathway in zebrafish, leading to disrupted fin formation in *ahr2* mutant fish. More investigation is required to determine at what stage and by what mechanism Ahr2 is involved in fin morphology and immune system function.

The increased severity of fin defects in our mutant compared to previously published mutants could suggest that previously published *ahr2* mutant zebrafish are hypomorphic rather than complete null alleles. Another possibility is that background mutations inherited along with *ahr2* mutations influence the fin phenotype. The presence of background mutations in the *ahr2^uab147/-^* line is unlikely, as heterozygotes were crossed to wild type for at least three generations before heterozygotes were crossed to each other to generate homozygous embryos. Strain differences could also account for phenotypic differences between mutants. *ahr2^hu3335^* and *ahr2^osu1^* mutants were generated on the 5D wild-type strain (Garcia, Bugel, Truong, Spagnoli and Tanguay, 2018; Goodale, La Du, Bisson, Janszen, Waters and Tanguay, 2012), while *ahr2^uab147^* mutants were generated on the AB wild-type strain. Regardless of the differences in severity of fin growth phenotype between different *ahr2* mutants, this study provides evidence that Ahr2 regulates fin maturation and/or maintenance independently of *ahr1* genes.

*ahr2* mutant larvae had grossly normal craniofacial structure while adults displayed craniofacial abnormalities. Previous studies using μCT imaging of *ahr2* mutant adult zebrafish reported abnormal skull morphology of several bones of the neurocranium, including the opercular bones, although morphology differences were not quantitated (Garcia, Bugel, Truong, Spagnoli and Tanguay, 2018; Goodale, La Du, Bisson, Janszen, Waters and Tanguay, 2012). In our analysis of *ahr* mutant skulls using μCT, we observed a narrowing of the ventral skull exclusively in *ahr2* mutants. This narrowed ventral skull shape in *ahr2* mutants is suggestive of disrupted skull patterning. Previous studies identified genes and pathways important for embryonic craniofacial development (Kimmel *et al.*, 2003; Kimmel *et al.*, 2007). However, the processes governing juvenile and adult craniofacial morphology are less well understood (Fisher *et al.*, 2003; Parsons *et al.*, 2011). Dermal skeletal development in zebrafish occurs during juvenile morphogenesis and adulthood to form structures including the lateral opercular bones, fin rays, and scales (Sire and Huysseune, 2007). The observation of severely deformed fins coupled with disproportionate ventral skull width in *ahr2* mutants suggests that Ahr2 may play a role in post-embryonic dermal skull development in zebrafish. In a study screening for mutants that affect post-embryonic zebrafish development, one mutation classified as “broadly affecting the dermal skeleton” (*dmh3* mutant) was identified as a missense mutation in the *edar* receptor (Henke *et al.*, 2017). This receptor severely disrupts fin growth, as described above, and was also shown to affect scale formation. Although skull morphology was not reported for this mutant, it is possible that the ventral skull defect observed in *ahr2* mutants is related to the *edar* pathway. Further work is required to support or reject this hypothesis, including analysis of scale development and morphology in adult *ahr2* mutants. Additionally, future work should determine the precise developmental stage at which fin and skull defects first appear.

Though the present study and previous work demonstrate roles for Ahr2 in fin and craniofacial development, less is known about the function of Ahr1a and Ahr1b in zebrafish. Further, ligands with the ability to activate Ahr1a or Ahr1b in vivo have not been characterized. A recent study evaluated the function *ahr* genes in vascular development in zebrafish embryos using an *ahr1a* single mutant, a double *ahr1b/ahr2* mutant and a triple *ahr1a/ahr1b/ahr2* mutant (Sugden, Leonardo-Mendonça, Acuña-Castroviejo and Siekmann, 2017). All mutants were grossly normal during development up to 5 dpf, consistent with normal morphology in our mutant embryos. When comparing endogenous and βNF-induced expression of the AHR target gene *cyp1a1*, they found that *ahr1a* mutants had normal expression patterns of *cyp1a1*, while *ahr1b/ahr2* and *ahr1a/ahr1b/ahr2* mutants lost both endogenous and βNF-induced *cyp1a1* expression. This is in contrast to *ahr2* single mutants, which retained partial endogenous and βNF-induced *cyp1a1* expression. These results support the theory that Ahr1b, but not Ahr1a, is an active AHR paralogue. It is possible, however, that Ahr1a may activate other target genes that were not studied. Further, no toxicity or abnormal morphology was reported with βNF treatment in wild-type embryos or in *ahr* mutant embryos. Therefore, the biological relevance of Ahr1b-specific *cyp1a1* activation is unclear. Other studies using morpholino knockdown of *ahr* genes have shown that Ahr1a, but not Ahr1b, contributes to the toxicity of PAHs and PCB-126 (Garner, Brown and Di Giulio, 2013; Incardona, Day, Collier and Scholz, 2006). Evidence from *in silico* molecular docking studies suggests that another AHR ligand, leflunomide, is capable of binding each of the zebrafish aryl hydrocarbon receptors with comparable binding affinities (Goodale, La Du, Bisson, Janszen, Waters and Tanguay, 2012; O’Donnell *et al.*, 2010). Further evidence in leflunomide-treated embryos suggests that Ahr1a induces expression of *cyp1a1* in the liver in the absence of both Ahr1b and Ahr2. Combined with our results that Ahr1a and Ahr1b are not required for TCDD-dependent toxicity in embryos, these results suggest that zebrafish ahr1 receptors are not required for canonical AHR signaling, but may have ligand- and tissue-specific effects outside of TCDD-mediated toxicity.

Though the previously cited studies, and the current study, suggest *ahr* genes are not essential for embryonic or larval development, this study is the first to characterize the effect of *ahr1a* and *ahr1b* mutation on adult morphology. In contrast to *ahr2* mutants, which display dysmorphic fins and skulls in adulthood, *ahr1* single and double mutants are morphologically normal. This suggests that *ahr2* is the primary zebrafish *AHR* paralogue regulating both endogenous and environmentally-induced AHR activity. We cannot rule out, however, that less obvious phenotypes are present in *ahr1* mutants that affect normal physiology or susceptibility to environmental stress.

Additionally, it should be noted that it is possible that the lack of phenotypes in *ahr1* mutants are due to hypomorphic alleles.

In addition to investigating the function of Ahr1a and Ahr1b in toxicity and development, we also investigated AHR/ER crosstalk in zebrafish embryos. AHR has been shown to regulate ER function both negatively and positively via direct protein-protein interaction and ubiquitination (Ohtake, Baba, Takada, Okada, Iwasaki, Miki, Takahashi, Kouzmenko, Nohara, Chiba, Fujii-Kuriyama and Kato, 2007; Ohtake, Takeyama, Matsumoto, Kitagawa, Yamamoto, Nohara, Tohyama, Krust, Mimura, Chambon, Yanagisawa, Fujii-Kuriyama and Kato, 2003). Specifically, ERα was recruited with AHR to the CYP1A promoter *in vitro* following TCDD treatment (Matthews, Wihlen, Thomsen and Gustafsson, 2005). If ERα is a necessary cofactor for TCDD-mediated AHR activity, inhibition or deletion of ERα should abrogate the response to AHR activation. We found, however, that pharmacologic or genetic inhibition of any of the zebrafish estrogen receptors failed to prevent Ahr2-dependent TCDD toxicity in embryos. Our results demonstrate that ERs are not constitutive partners of AHR and supports the hypothesis that AHR cofactor recruitment is context dependent.

## ACKNOWLEDGLEMENTS

We thank Jeffery King and Marijke Schrock for technical support. We thank Dr. Susan Farmer and Angela Mannone and the fish caretakers for help with zebrafish husbandry. We thank Dr. Matthew P. Harris for advice on micro-CT analysis. This work was supported by National Institute of Environmental Health Sciences (R01ES026337 to DAG) and National Institute of General Medical Sciences (T32GM008361 to JPS).

